# High-resolution CRISPR/Cas9 screens identify PAK2 as a suppressor of macrophage proliferation

**DOI:** 10.64898/2026.09.14.751564

**Authors:** Kevin N. Wanniarachchi, Astha Neupane, Bidhi Kasu, Bijaya Pradhan, Susmita Poudel, Joshua J. Balster, Jason G. Kerkvliet, Adam D. Hoppe, Natalie W. Thiex

## Abstract

Macrophage abundance is regulated by pathways controlling survival and proliferation, yet the genetic determinants of macrophage fitness remain poorly understood. Here, we performed replicate genome-wide CRISPR/Cas9 screens in primary murine bone marrow-derived macrophages (BMDMs) to identify positive and negative regulators of macrophage fitness. Validity of the screens is supported by finding previously characterized core essential genes such as those encoding translation machinery, while disruption of established tumor suppressors increased BMDM representation in the culture. As expected, *Csf1r* encoding the macrophage growth factor emerged as a positive regulator of macrophage fitness. Unexpectedly, *Pak2*, encoding p21-activated kinase 2 (PAK2), emerged as a negative regulator of macrophage fitness. Targeted *Pak2* disruption increased BMDM accumulation and was associated with elevated cyclin-D1 expression, identifying PAK2 as a suppressor of macrophage proliferation. PAK2 also localized to CSF1-induced actin-rich membrane ruffles, yet its depletion did not impair ruffle formation. Instead, PAK2-depleted macrophages exhibited persistent F-actin-rich ruffles, larger macropinosomes, and increased fluid-phase uptake. These changes were accompanied by altered LIMK/cofilin signaling and persistent cofilin localization at macropinocytic structures. Phosphoproteomic analysis further identified reduced phosphorylation of proteins associated with cytoskeletal regulation, phosphoinositide signaling, and endosomal trafficking. Together, these findings identify PAK2 as a context-dependent regulator that restrains macrophage proliferation and membrane-remodeling activity and demonstrate that PAK2 has markedly different fitness functions in primary macrophages than previously identified in several transformed cell types.

## Introduction

Macrophages are highly specialized immune cells that contribute to tissue homeostasis, host defense, repair, and the pathogenesis of inflammatory, cardiovascular, and neoplastic diseases (Moore and Tabas, 2011; Wynn *et al*., 2013; Wynn and Vannella, 2016). Their abundance within tissues is determined not only by recruitment and differentiation from circulating progenitors, but also by survival and local proliferation of differentiated macrophages (Gentek *et al*., 2014; Blériot *et al*., 2020; Gessain *et al*., 2020). These properties make the mechanisms controlling macrophage fitness, which is the capacity of cells to survive, proliferate, and persist under a given environmental condition (Hart *et al*., 2015), important determinants of macrophage function in both homeostasis and disease. Defining the pathways that control macrophage fitness is also increasingly relevant to therapeutic strategies that seek to manipulate macrophage function or employ engineered macrophages as cell-based therapies for cancer, inflammatory disease, and tissue repair (Mantovani *et al*., 2017; Sergin *et al*., 2017; Klichinsky *et al*., 2020; Na *et al*., 2023; Zhang *et al*., 2023).

Macrophages have specialized cellular demands that may create distinct requirements for survival and proliferation. Unlike rapidly proliferating transformed cells commonly used in genome-wide fitness studies, differentiated macrophages must maintain long-term viability while supporting extensive cytoskeletal remodeling, membrane trafficking, phagocytosis, endocytosis, and growth-factor-dependent signaling (Swanson, 2008). These processes place unusual demands on pathways controlling metabolism, organelle function, membrane dynamics, and cell-cycle regulation (Doodnauth *et al*., 2019). Consequently, genes that promote fitness in cancer or other proliferative cell types may have different, or even opposing, functions in macrophages, emphasizing the need to define fitness regulators directly in primary macrophages. Although genome-wide fitness screens have defined essential and context-dependent dependencies in numerous cancer models (Hart et al., 2015; Behan et al., 2019), extrapolating these relationships to primary macrophages may overlook lineage-specific mechanisms controlling cell abundance, survival and even adherence.

Genome-wide CRISPR/Cas9 screening provides an opportunity to directly define these dependencies by systematically disrupting genes within a defined cellular context and has provided resolution beyond essential genes to identify fitness genes at high resolution in mammalian systems (Ramani *et al*., 2012; Hart *et al*., 2015; Wang *et al*., 2015). In the context of cancer, identifying fitness genes can reveal cell type-specific vulnerabilities that can be targeted therapeutically (Tzelepis *et al*., 2016; Chen *et al*., 2017; Behan *et al*., 2019) A whole-genome CRISPR screen in immortalized bone marrow-derived macrophages identified genes required for macrophage viability and inflammatory signaling, demonstrating the utility of this approach for interrogating macrophage biology (Covarrubias *et al*., 2020). However, immortalization itself alters growth-control pathways, making it important to determine fitness requirements in primary macrophages (Andreu *et al*., 2017). We therefore performed replicate genome-wide CRISPR/Cas9 screens in primary bone marrow-derived macrophages (BMDMs), introducing gene disruptions after the cells had undergone several days of CSF1-dependent differentiation and became adherent.

Our screens identified both positive and negative regulators of macrophage fitness. Genes whose disruption reduced BMDM abundance included broadly required cellular fitness genes involved in core processes such as proteostasis and translation (e.g., *Hspa5* and *Rps6*), as well as canonical macrophage-associated genes including *Csf1r* and *Itgam* (CD11B). Conversely, disruption of established tumor suppressors and growth regulators, including *Trp53* (P53) and *Pten* (PTEN), increased BMDM abundance, validating the ability of the screens to identify negative regulators of macrophage fitness. The screens also identified genes whose fitness phenotypes may reflect macrophage-selective or strongly context-dependent functions. Among these, *Pak2*, encoding p21-activated kinase 2 (PAK2) (Renkema *et al*., 2002), emerged as a key negative regulator of macrophage fitness despite class I PAKs being classified as an essential and generally supporting cellular fitness in several other contexts (Buikhuisen *et al*., 2023)

## Methods

### Reagents

Bone marrow-derived macrophages (BMDM) were cultured in bone-marrow medium (BMM) containing 50% high glucose Dulbecco’s Modified Eagle Medium (ATCC 30-2002, American Type Culture Collection, Manassas, VA), 20% heat-inactivated fetal bovine serum (Atlanta Biologicals), and 30% L-cell supernatant (Stanley and Heard, 1977) a source of colony-stimulating factor-1, 10,000 IU penicillin and 10 mg/mL streptomycin (Corning, Manassas, VA), and 5.7 mM 2-mercaptoethanol. DPBS without calcium or magnesium was purchased from GE Healthcare Life Sciences, Pittsburgh. Cyclosporin A (30024, Sigma). Mouse Brie CRISPR knockout pooled library developed by David Root and John Doench was obtained from Addgene (Addgene #73633) (Doench *et al*., 2016)

**Table 1.** Antibodies.

| Target Protein | Species | Isotype | Mono/Poly | Manufacturer | Catalog number |
| --- | --- | --- | --- | --- | --- |
| 4E-BP1 | Rabbit | not listed | Polyclonal | CST | 9452 |
| Beta actin | Mouse | IgG2b | Monoclonal | CST | 3700 |
| Cofilin | Mouse | IgG1-Kappa | Monoclonal | Santa Cruz | 376476 |
| GAPDH | Mouse | IgG1-Kappa | Monoclonal | Santa Cruz | 47724 |
| LIMK1 | Mouse | IgG1-Kappa | Monoclonal | Santa Cruz | 515585 |
| MTOR | Rabbit | IgG | Monoclonal | CST | 2983 |
| NF2 | Mouse | IgG1-Kappa | Monoclonal | Santa Cruz | 55575 |
| p21 WAF1/CIP1 | Mouse | IgG2b-kappa | Monoclonal | Santa Cruz | 6246 |
| PAK2 | Rabbit | IgG | Monoclonal | CST | 2615 |
| Phospho-4E-BP1 (Thr37/46) | Rabbit | IgG | Monoclonal | CST | 2855 |
| Phospho-70 S6 kinase (Ser371) | Rabbit | not listed | Monoclonal | CST | 9208 |
| Phospho-70 S6 kinase (Thr389) | Rabbit | not listed | Monoclonal | CST | 9234 |
| Phospho-Cofilin (Ser3) | Rabbit | IgG | Monoclonal | CST | 3313 |
| Phospho-MTOR (Ser2448) | Rabbit | IgG | Monoclonal | CST | 5536 |
| Phospho-NF2 | Rabbit | not listed | Polyclonal | CST | 9163 |

### Brie library lentivirus production for CRISPR/Cas9 whole genome screens

Brie library amplification, lentiviral production, and functional titer determination were performed as described (Joung *et al*., 2017) with minor modifications. Briefly, the Brie library plasmid DNA was amplified in Stable3 competent *E. coli* and subjected to next-generation sequencing (NGS) to determine the distribution of sgRNAs in the library (Input). Approximately 1.8 × 10^6^ HEK 293T cells were seeded per 100mm plate and were transfected the following day with 6 µg sgRNA library plasmids (Addgene# 73633), 6 µg psPAX2 plasmid, 1 µg VSV-G plasmid, and 24 µg polyethyleneimine (PEI). The functional titer of the lentivirus was determined by live cell counting after puromycin selection.

### Bone marrow-derived macrophage (BMDM) isolation and culture for CRISPR/Cas9 screens

Transgenic mice with a Rosa26-Cas9 knock-in on a C57BL/6J background were received from Jackson Laboratories (Stock No. 026179, Bar Harbor, ME). Mice were euthanized using CO_2_ inhalation and cervical dislocation, femurs were dissected from the mice, and bone marrow was harvested (Racoosin and Swanson, 1989) (Swanson, 1989) by flushing Dulbecco’s phosphate buffer saline (DPBS) without calcium or magnesium (GE Healthcare Life Sciences, Pittsburgh, PA) through the bone using a needle and syringe. Cells were plated in culture media containing 20% HI-FBS and 30% L-cell supernatant in high glucose DMEM in non-tissue culture-treated sterile dishes and incubated at 37°C with 5% CO_2_. Additional culture media was added to the cells on day two post-isolation. Following cell adherence to the dish, on day 4, the medium was removed entirely, and fresh BMM was added to the culture. On day five cells were treated with 10 mM Cyclosporin A for 20 minutes before adding lentivirus with Brie library to the cells. Multiplicity of infection (MOI) ∼ 0.3-0.1. Library coverage was around 100-200-fold representation of each sgRNA. Transduced cells were antibiotic-selected two days after transduction with 5mg/mL puromycin. Cells were cultured for at least 14 days for protein depletion (post-culture).

### DNA isolation and next-generation sequencing

Genomic DNA was extracted from post-culture cells using the GeneJET genomic DNA extraction kit (#K0721 ThermoFisher). The sgRNA inserts from genomic DNA (Post-culture) and library plasmid DNA (Input) were PCR amplified by a single-step PCR protocol (Broad Institute). Barcoded primers with staggered sequences are used for next-generation sequencing (NGS). Sequencing of 75 base reads was conducted on Illumina Nextseq 500. Following quality control analysis using FASTQC, reads were mapped to the sgRNA library to identify genes and read frequency tables were generated using Model-based Analysis of Genome-wide CRISPR-Cas9 Knockouts (MAGeCK), version 0.5.9 (Li *et al*., 2014).

### Statistical Analysis of Sequencing Data and Gene Ranking

MAGeCK version 0.5.9 was used to generate read frequency tables of all sgRNAs and map sgRNAs to the Brie library to identify corresponding genes with no mismatches tolerated. Specifically, the count function (-count) in MAGeCK was used to generate frequency tables for each population. Next, we used MAGeCK-RRA using the paired test function (-test -paired) to compare sgRNA read frequencies between input and post-culture populations. This allowed for the comparison of replicate screens. Additionally, the Brie library contains ∼1000 control non-coding sgRNA inserts. We considered genes with a false discovery rate of equal or less than 0.01 as cutoff criteria for significance for MAGeCK-RRA analysis.

### Pathway Enrichment Analysis

Ranked lists of genes from the paired analysis of screens were used to identify biologically enriched pathways using the web-based tools iDEP.96 (Integrated Differential Expression and Pathway Analysis (Ge *et al*., 2018). Briefly, we filtered the top 100 ranked genes based on MAGeCK rank based on RRA-score, and then this ranked list with each gene log-fold change value was uploaded to iDEP.96. We used the Gene Set Enrichment Analysis (GSEA) to analyze the pathway enriched within the ranked lists.

### Targeted sgRNA transductions to validate genes

To make targeted sgRNA-mediated gene disruptions, sgRNAs from the Brie library were selected and cloned to lenti-guide-puro plasmid (Addgene #52963). SgRNA binding sites on the mouse genome were confirmed using the NCBI blast tool (Supplemental Figure 1) (Altschul *et al*., 1990). Plasmids were amplified in Stable3 competent *E. coli*. Purified plasmid DNA was used to transfect HEK293T cells described before to produce sgRNA containing lentivirus. Viral supernatant with 10 µM Cyclosporin A was added to Cas9-expressing BMDMs cultured in BMM for 48 h and was selected with puromycin. Validation assays were done 16 days post-transduction.

**Figure 1.**
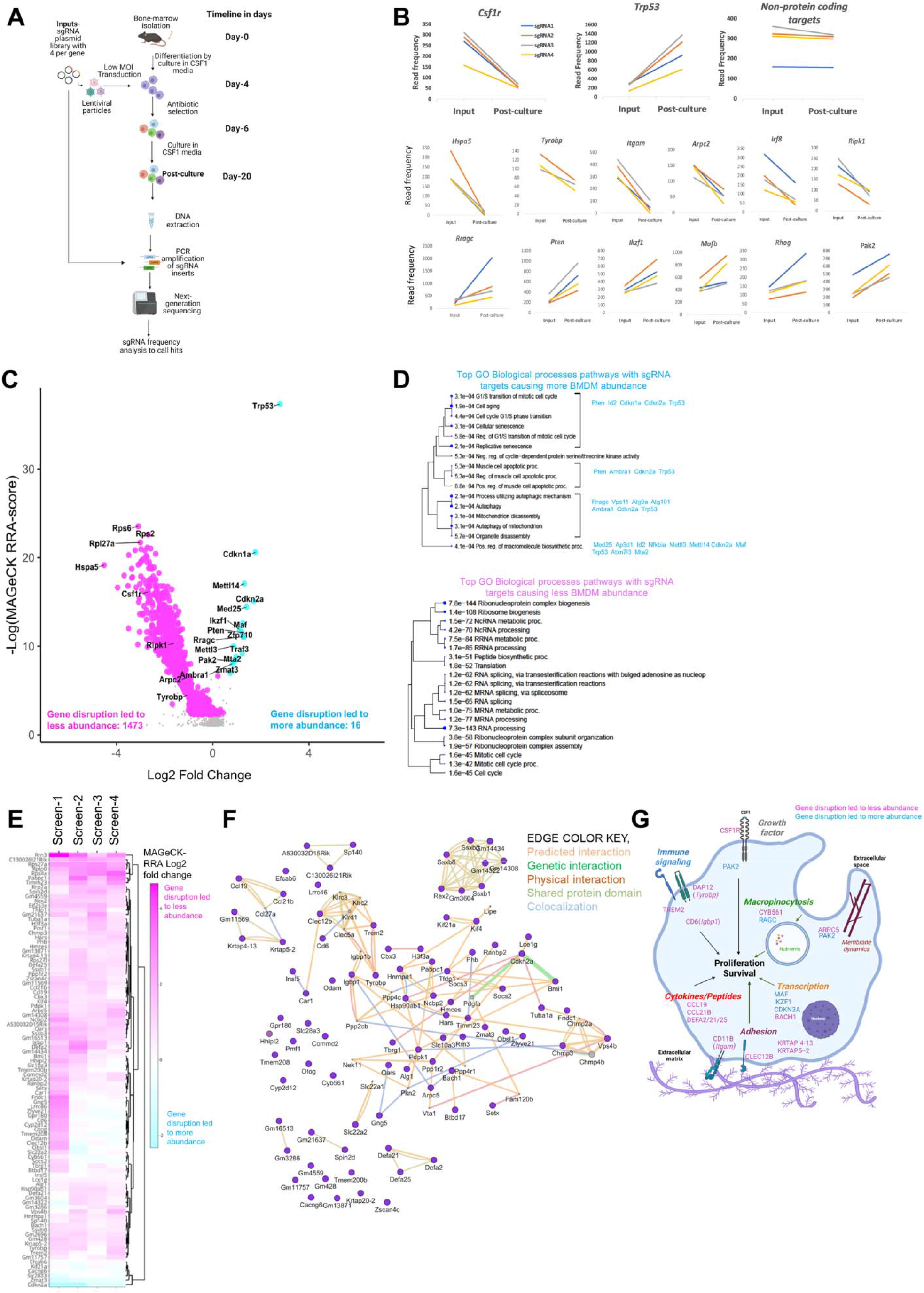
Whole genome CRISPR/Cas9 screens identify sgRNAs that resulted in a decrease or increase in cell abundance of BMDMs in culture. A. Bone marrow cells from mice were isolated and cultured in CSF-1-rich media for four days post-isolation. Subsequently, the cells were transduced with the sgRNA at low MOI (0.2-0.3), which included four sgRNA per gene targeting approximately 19,000 protein-coding genes in the mouse genome. Following the transduction, antibiotic selection was done, and cells were cultured for about 12-14 days in a CSF1-containing medium. Next-generation sequencing was performed to determine the frequency of sgRNA in both the plasmid library and post-culture populations. B. sgRNA read counts from a single screen of selected genes such as macrophage essential gene Csf1r shows gene disruption leading to lower abundance, sgRNA targeting tumor suppressor *Trp53* (P53) led to higher abundance in the post-culture pool. sgRNA targeting non-protein coding regions (4 different regions) did not affect abundance. Targeting *Hspa5, Tyrobp* (DAP12), *Itgam* (CD11B), *Arpc2, Irf8, Ripk2* led to lower abundance. Targeting *Rragc, Pten, Ikzf1, Mafb, Rhog*, and *Pak2* led to higher abundance. X-axis read count, y-axis Input plasmid DNA, and post-culture pool. C. MAGeCK-RRA identifies macrophage fitness genes. sgRNA read counts from 4 independent screens were analyzed by the MAGeCK-RRA algorithm to identify genes. Data is cumulative MAGeCK-RRA paired analysis of sgRNA read counts from 4 independent screens. Significance cutoff of colored genes FDR<=0.005 D. Biological pathways enriched within the top 100 ranked genes were identified by gene-set enrichment analysis (GSEA). Bar plots with the number of genes in each pathway via the ShinyGO tool. E. Unique BMDM fitness genes were identified by comparison with the human essential gene list (DEG15). 95 out of 1283 of our gene list was unique, suggesting importance in BMDM fitness. Heatmap shows MAGeCK-RRA LFC of these 95 genes across individual screens. F. Interaction analysis of unique macrophage fitness genes. Edges represent associated interaction or protein domain similarity between two gene products based on literature. Generated using GeneMania on Cytoscape. G. Model showing the cellular function of selected screen hits.

### BMDM proliferation assay and bicinchoninic acid assay (BCA)

Post-transduction day 16 BMDM were plated at equal density on three separate 24-well culture plates. Here, the culture media was prepared Bone marrow medium (BMM) containing 20% premium heat-inactivated fetal bovine serum (FBS) (Atlanta Biologicals, Flowery Branch, GA), 50ng/ml recombinant mouse CSF1 (BioLegend), 10,000 I.U penicillin and 10 mg/mL streptomycin (Corning, Manassas, VA), 5.7 mM 2-mercaptoethanol, and Dulbecco’s Modified Eagles Medium (DMEM) containing 4.5 g/L glucose, 0.5 g/L L-glutamine and 0.1 g/L sodium pyruvate. On the day of seeding (day 0), day 2, and day 4, a single 24-well plate was imaged following live-cell nuclei staining. For each genotype, nine fields of view across three technical replicates were imaged per experiment. Following imaging, a BCA assay was conducted. Cells were lysed with cold lysis buffer (M-PER mammalian protein extraction reagent (78503, ThermoFisher Scientific, Waltham, MA). BCA assay was done using the Pierce BCA protein assay kit (23225, ThermoFisher Scientific, Waltham, MA), and following the BCA reaction, plate intensity was measured using a spectrophotometer.

### Cut-site sequencing to confirm gene disruption

Genome editing efficiency was verified by targeted amplicon sequencing of CRISPR cut sites. Between 0.5 × 10^6 and 2 × 10^6 puromycin-selected BMDMs were harvested 7–10 days after lentiviral transduction and pelleted by centrifugation. Cell pellets were washed with PBS, frozen at −80°C, and genomic DNA was isolated using the GeneJET Genomic DNA Purification Kit. PCR primers flanking each sgRNA target site were designed using CHOPCHOP software with an expected amplicon size of 350–450 bp. Target loci were amplified using Phire Hot Start II DNA Polymerase in 50-μL reactions containing 0.3 μM forward and reverse primers and up to 1 μg genomic DNA. PCR products were purified using AMPure XP magnetic beads, quantified, and submitted to Plasmidsaurus for nanopore sequencing using the Premium PCR sequencing service. Sequencing data were analyzed using CRISPResso2 (Clement *et al*., 2019) to determine insertion/deletion frequencies and the proportion of remaining wild-type alleles. Editing efficiencies were calculated from the percentage of modified alleles relative to total aligned sequencing reads.

### Immunofluorescence staining

BMDM were plated on glass coverslips and were fixed with 4% PFA in 1XPBS for 12 mins. Permeabilized in 0.3% Triton-X for one hour at room temperature and blocked in 5% bovine serum albumin for one hour. Primary antibodies were diluted in 1% BSA in PBS before being added to cells overnight and incubated at 4°C. Appropriate fluorescent secondary antibodies against primary antibodies were used. Nuclei were stained with DAPI. Images were acquired on a wide-field epifluorescence Olympus IX83 microscope using 40X air objective and 60X oil objective. Confocal images were acquired using a Leica spinning disk microscope with a 60X oil objective. Image prepared for montages using ImageJ.

### Cell lysate preparation and SDS PAGE

Cell lysates were prepared by incubating the cells in lysis buffer (M-PER mammalian protein extraction reagent (78503, ThermoFisher Scientific, Waltham, MA) supplemented with Halt protease inhibitor cocktail (1:50, ThermoFisher Scientific, Waltham, MA) and phosphatase inhibitor cocktail (1:100, ThermoFisher Scientific, Waltham, MA) and 10 mM sodium orthovanadate for 10 min at 4°C with cell scraping to make sure complete lysis. Lysates were harvested by centrifuging at 18,000 g for 15 min. Protein concentration was determined by performing a BCA assay using the Pierce BCA protein assay kit (23225, ThermoFisher Scientific, Waltham, MA). 20 µg total protein was loaded per well of 4-20% SDS-PAGE precast gel. Proteins were transferred to the Polyvinylidene fluoride (PVDF) membrane via the wet transfer in 20% MeOH. PVDF membranes were blocked in 5% non-fat milk prepared in 1x Tris-buffered saline with Tween 20. Primary antibodies were diluted as recommended by the manufacturer and incubated overnight at 4°C. Membranes were imaged using LICOR Fc Odyssey, and band intensities were analyzed on Image Studio Lite software.

### RNA isolation and RT-qPCR

Cells were lysed to extract RNA using the RNeasy Mini Kit from Qiagen. Residual DNA was digested using DNAse-I from ThermoFisher. For RT-qPCR, 50ng RNA was prepared as recommended for the iTaq Universal SYBR Green One-step kit from BioRad. qPCR reactions were prepared in triplicate for target mRNA. Ubiquitin-c was used as a reference control for qPCR in triplicate per sample. Amplification was as follows: reverse transcription reaction 10min at 50°C, polymerase activation, DNA denaturation 1 min at 95°C, amplification cycles of 15-second denaturation, and 60-second annealing/extension for 45 cycles. Cq was calculated using Design and Analysis 2.6.0.

### Data availability

Next-generation sequencing output, processed read count frequency tables, and MAGeCK test output are available under GEO accession GSE251887.

## Results

### High-resolution CRISPR/Cas9 whole genome fitness screens identify genes regulating the proliferation and survival of BMDM

To identify genes regulating bone marrow-derived macrophage (BMDM) fitness, we used the Brie pooled sgRNA library to transduce day-4 post-isolation BMDMs and generate gene-disrupted populations, which were subsequently antibiotic selected and cultured for 12-14 days (Figure 1A). Because sgRNA delivery occurred after several days of CSF1-dependent culture and cell adherence, this strategy was designed to identify genes that regulate the subsequent survival and proliferation of nascent macrophages rather than genes required for macrophage differentiation. We reasoned that disruption of genes that support BMDM fitness would reduce the relative abundance of the corresponding sgRNAs during the post-transduction culture period, whereas disruption of genes that normally restrict macrophage proliferation or survival would increase their relative abundance. To quantify sgRNA representation, lentiviral sgRNA inserts were amplified from genomic DNA isolated from the post-culture BMDM population and analyzed by next-generation sequencing. The sgRNA abundance in the post-culture population was compared with the input lentiviral plasmid library using MAGeCK (Li *et al*., 2014) for four independent screens to generate robust rank aggregation (RRA) scores, gene ranks, and false discovery rates (FDR) (Supplemental Table 1).

The screen recovered expected regulators of macrophage fitness and identified novel hits. For example, sgRNAs targeting *Csf1r*, which encodes the CSF1 receptor required for macrophage survival and proliferation, were strongly depleted from the post-culture population, whereas sgRNAs targeting the tumor suppressor *Trp53* were strongly enriched (Figure 1B). In contrast, sgRNAs targeting non-protein-coding control regions remained equally distributed (Figure 1B).

Next, we highlighted representative hits from one of the replicate screens to illustrate novel macrophage-specific fitness genes. Genes that decreased in abundance in individual screens included *Hspa5*, an ER chaperone required for proteostasis (Lee, 2001); *Tyrobp* and *Itgam*, which participate in macrophage receptor signaling, adhesion, and phagocytic functions (Fagerholm *et al*., 2013; Rotty *et al*., 2017; Haure-Mirande *et al*., 2022); *Arpc2*, a component of the Arp2/3 actin-nucleating complex (Rotty *et al*., 2017); *Irf8*, a key regulator of myeloid cell differentiation and identity (Xia *et al*., 2020); and *Ripk2*, a kinase linking NOD-family receptors to innate immune signaling (Rivoal *et al*., 2023) (Figure 1B). In contrast, disruption of *Rragc, Pten, Ikzf1, Mafb, Rhog*, and *Pak2* increased BMDM abundance in representative screens, implicating regulators of nutrient sensing and mTOR signaling, PI3K signaling, hematopoietic transcriptional programs, and Rho-family/cytoskeletal signaling as potential negative regulators of macrophage fitness (Sancak *et al*., 2008; Aziz *et al*., 2009; Tzircotis *et al*., 2011). Together, the recovery of genes spanning macrophage identity, innate immune signaling, cytoskeletal organization, and growth-control pathways illustrates the ability of these screens to identify both established and potentially macrophage-selective determinants of fitness.

Using paired MAGeCK-RRA analysis to generate a composite ranking of genes that reproducibly altered BMDM abundance from the four replicate screens, we identified 1,473 genes whose disruption decreased representation in the post-culture population and 16 genes whose disruption increased representation, defining a broad set of candidate positive and negative regulators of macrophage fitness (Figure 1C, Supplemental Table 1).

Gene set enrichment analysis (GSEA) showed that genes whose disruption decreased BMDM abundance were strongly enriched for core cellular processes, including ribosome biogenesis, RNA processing and splicing, translation, DNA replication, and cell-cycle progression (Figure 1D). In contrast, genes whose disruption increased BMDM abundance were enriched for pathways associated with cell-cycle control, senescence, apoptosis, autophagy, and organelle turnover, consistent with the identification of genes that normally restrict macrophage proliferation or survival (Figure 1D).

The Database of Essential Genes offers a comprehensive list of human essential genes (Luo, 2021). Around 92.6% of the significant genes identified by our screens overlapped with the DEG list, indicating that those genes are critical in fundamental biological processes across many or all cells, while 7.4% of genes (95 genes) identified here are unique (Supplemental Table 2). We extracted the 95 unique macrophage fitness genes and displayed the MAGeCK RRA Log2-fold change from the replicate screens as a heatmap to visualize consensus among individual replicate screens and to facilitate generation of future macrophage-specific fitness hypotheses (Figure 1E). Of note, *Pak2* is not included among the 95 BMDM-unique genes because it is represented in the Database of Essential Genes as an essential gene, whereas our primary macrophage screens have identified it as a suppressor of macrophage proliferation.

To determine reported interactions between the proteins encoded by these unique macrophage genes, we utilized the protein-protein interaction analysis tool Genemania (Figure 1F) (Warde-Farley et al., 2010).

Based on the interaction map generated, known interactions such as immune receptor signaling adaptor *Tyrobp* (DAP12) and immune cell receptor TREM2 were both enriched within the macrophage unique genes list. Several of these selected genes and the encoded protein localization based on previous studies are shown in the diagram (Figure 1G). Together, these analyses distinguish broadly conserved cellular dependencies from genes and pathways whose effects on fitness may be particularly context-dependent in macrophages.

### PAK2 limits macrophage proliferation and Cyclin D1 expression

*Pak2* was identified in the fitness screens as a negative regulator of BMDM abundance. To validate this phenotype, PAK2-depleted BMDMs were generated using sgRNAs targeting *Pak2*. Efficient depletion of PAK2 protein was confirmed by immunoblotting, and editing at the targeted *Pak2* locus was independently verified by cut-site sequencing (Figure 2A; Supplemental Figure 2). Equal numbers of control and PAK2-depleted BMDMs were subsequently plated and cultured in CSF1-containing medium. PAK2-depleted BMDMs increased in abundance over four days relative to control cells, consistent with enhanced proliferation following loss of PAK2 (Figure 2B). Increased cell accumulation was independently supported by higher total protein concentration in PAK2-depleted cultures at day 4 (Figure 2C).

**Figure 2.**
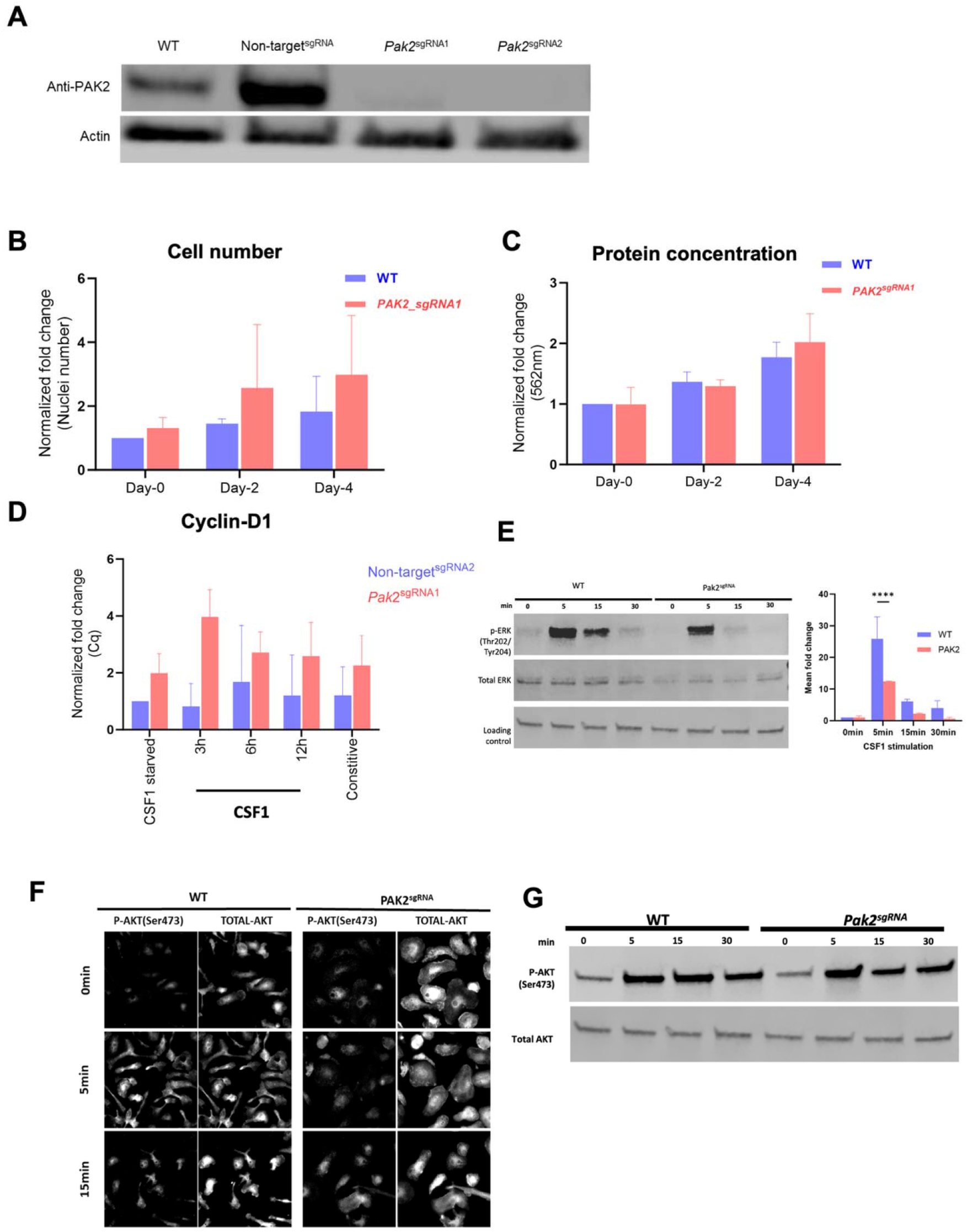
PAK2 is a suppressor of macrophage fitness. A. Protein depletion following transduction with sgRNA targeting validated through western blot. Day-16 post-transduction cells were blotted with anti-PAK2 antibody. B. *Pak2*^sgRNA^ gene disruption led to an increased abundance of BMDMs in culture relative to control non-target^sgRNA^ following a 4-day culture period. After protein depletion, an equal density of cells were seeded and cultured in media containing 50 ng/mL recombinant CSF1. Nuclei were stained and imaged. In each independent experiment, nine fields of view across three technical replicates were imaged. The bar graph shows normalized fold change across three independent experiments. Error bars are the standard deviation of fold change. ***adjusted p-value<0.0001. C. Culture wells with *Pak2*^sgRNA^ BMDM had higher protein concentration than control non-target^sgRNA^ BMDM. An equal density of cells was seeded and cultured in media containing 50 ng/mL recombinant murine CSF1. On indicated days, cells were lysed, and protein concentration was measured following the bicinchoninic acid reaction using a spectrophotometer. In each independent experiment, three technical replicate wells were measured. The bar graph shows normalized fold change across three independent experiments. Error bars are the standard deviation of fold change. **adjusted p-value<0.002 D. PAK2 depletion causes an increase in cyclin-D1 levels independent of CSF1 stimulation. Control non-target^sgRNA^ cyclin-D1 levels increased with time following 3-12 h of CSF1 stimulation. Comparatively, *Pak2*^sgRNA^ had higher levels of cyclin-D1 at all time points. mRNA were quantified in post-isolation day 20/post-transduction day 16 BMDM by one-step RT-PCR. E. Activation of ERK1/2 by phosphorylation following CSF1 stimulation is lowered targeted gene disrupted BMDM. Here BMDMs were starved of CSF1 overnight and were incubated with 200 ng/mL rmCSF1 for indicated times. Bar graph showing ratio of total to phosphorylated ERK from multiple experiment quantification of western blot band intensities. F. CSF1 stimulated AKT activation in is not altered in *Pak2*^*sgRNA*^ BMDM. Fluorescent images of AKT activation (Ser473) by phosphorylation following CSF1 stimulation in WT and gene disrupted BMDM. Here BMDMs were starved of CSF1 overnight and were incubated with 200ng/mL recombinant CSF1 for indicated times. Following stimulation BMDMs were 4% PFA fixed and prepared for anti-body labeling. Genotype images are window-leveled to WT 5min for each p-AKT and total AKT. G. Similar observation was made via western blot which showed relative to WT *Pak2*^*sgRNA*^ had similar level of p-AKT (Ser473)

Cyclin D1 mRNA was also elevated in PAK2-depleted BMDMs during a CSF1 stimulation time course, with higher expression observed relative to control cells across the measured time points (Figure 2D). This increase in cyclin D1 expression is consistent with the enhanced proliferative phenotype of PAK2-depleted macrophages. In contrast to other cell types in which PAK2 activity has been associated with maintenance of cyclin D1 expression and proliferation, these results indicate that PAK2 restrains cyclin D1 expression and cell proliferation in BMDMs.

Both ERK and AKT have well-characterized dynamics downstream of CSF1R stimulation. In an experimental model where cells are starved overnight of CSF1 and subsequently stimulated with CSF1, levels of phosphorylated AKT and ERK rise within minutes and then go back down (Huang *et al*., 2024) Therefore, we predicted excess proliferation in *Pak2*^*sgRNA*^ BMDM might correlated with increased Akt or Erk phosphorylation. However, neither ERK nor AKT activity was elevated or prolonged following CSF1 stimulation relative to WT BMDMs (Figure 2E-G).

### PAK2 localizes to dorsal membrane ruffles and limits macropinocytosis

PAK2 is a group I p21-activated kinase that functions downstream of Rho-family GTPases involved in actin remodeling, including Rac and Cdc42 (Manser *et al*., 1994; Hoppe and Swanson, 2004). PAK2 is also rapidly activated in macrophages following CSF1 stimulation (Weiss-Haljiti *et al*., 2004). To determine whether PAK2 localizes to membrane structures associated with CSF1-stimulated macropinocytosis, we examined endogenous PAK2 and F-actin in BMDMs following growth-factor stimulation by confocal microscopy. PAK2 was enriched along F-actin-containing membrane ruffles and circular dorsal ruffles within minutes of CSF1 stimulation (Figure 3A). PI3K activity is dispensable for the initial formation and morphology of macrophage membrane ruffles but is required for efficient closure of ruffles into macropinosomes (Quinn *et al*., 2021).

**Figure 3.**
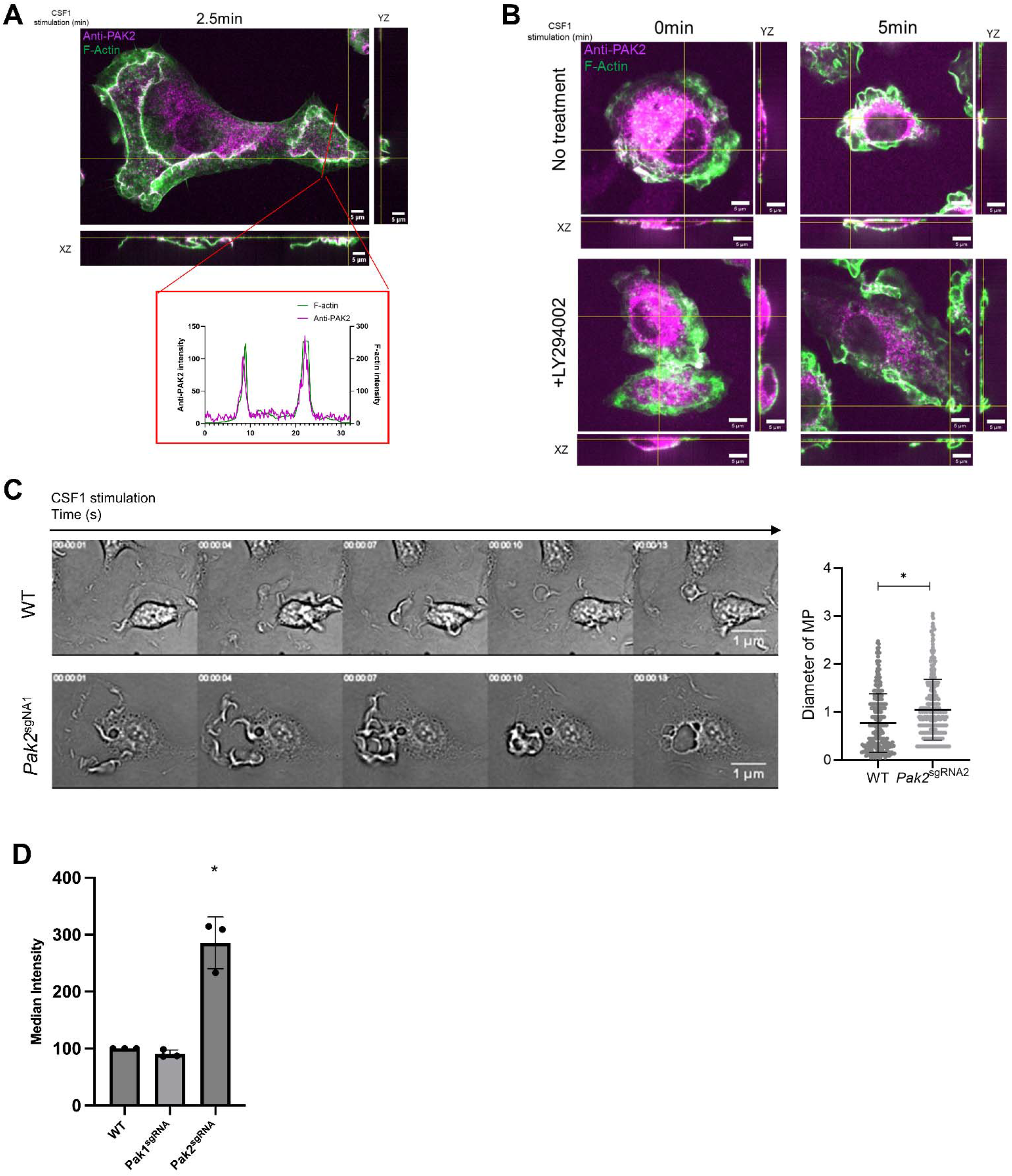
PAK2 localizes to circular dorsal ruffles and inhibits macropinocytosis. A. PAK2 (magenta) localizes to filamentous actin (Phalloidin-488 in green) in forming circular dorsal ruffles and laterally extending f-actin in the membrane following CSF-1 stimulation in BMDM. Confocal microscopy of WT BMDM starved of CSF1 overnight and stimulated with 200ng/mL CSF1 for 2.5min. B. PAK2 recruitment is independent of PI3K activity. PAK2 was seen along with f-actin-rich lamella following stimulation in both non-treated and PI3K-inhibited (with LY294002). Here, BMDM were starved of CSF1 overnight and incubated with LY294002 for 30 minutes before being stimulated with 200 ng/mL rec. Mouse CSF1 for indicated times. Each panel window-leveled to each other. C. *Pak2*^sgRNA^ BMDM displays more peripheral membrane ruffles and the formation of larger macropinosomes following stimulation with or without CSF1 stimulation. Suggesting that PAK2 has a role in membrane ruffle assembly and/or disassembly. BMDMs were starved of CSF1 O/N, stimulated with 200ng/mL CSF1, and imaged for 5 minutes (Supplemental Movie 1). The size of macropinosomes was measured by calculating ferret diameter using ImageJ as stated in the methods. D. Pak1 disruption does not phenocopy Pak2 disruption, suggesting that they do not play a redundant role in regulation macropinocytosis under CSF1-stimulated conditions.

Consistent with this distinction, LY294002-treated BMDMs continued to form prominent F-actin-rich ruffles after CSF1 stimulation, and PAK2 remained detectable at these structures (Figure 3B). Although these qualitative observations do not establish whether PI3K activity alters the extent of PAK2 recruitment, they place PAK2 within the actin-rich membrane domains that generate macropinosomes.

Disruption of *Pak2* produced a striking increase in membrane ruffling and generated significantly larger macropinosomes compared to WT BMDM following CSF1 stimulation (Figure 3C; Supplemental Movies 1-3), indicating that PAK2 is not required for ruffle formation or macropinosome generation in BMDMs but instead acts to restrict these processes. We next measured fluid-phase uptake using Lucifer yellow, which preferentially reports nonspecific fluid uptake in BMDMs rather than dextran where the MRC1-receptor-mediated uptake contributes substantially to internalization (Swanson *et al*., 1985; Wollman *et al*., 2024).

*Pak2*^sgRNA^ BMDMs exhibited increased Lucifer yellow uptake relative to control cells, whereas disruption of the closely related group I kinase *Pak1* did not have any effect on Lucifer yellow uptake (Figure 3D). These results identify a nonredundant role for PAK2 in limiting macrophage macropinocytosis. Because macropinocytosis can provide extracellular nutrients and modulate nutrient-sensitive signaling pathways in macrophages, enhanced macropinocytic activity following PAK2 depletion could contribute to changes in cellular metabolic or growth state that drive the proliferative phenotype (Yoshida *et al*., 2015)

### PAK2 regulates actin turnover and LIMK/cofilin signaling during CSF1-induced membrane ruffling

Both WT and *Pak2*^sgRNA^ BMDMs formed prominent F-actin-rich membrane ruffles rapidly following CSF1 stimulation, indicating that PAK2 is not required for the initial actin polymerization and ruffling response. In WT BMDMs, these structures were transient, whereas F-actin-rich ruffles persisted in *Pak2*^sgRNA^ BMDMs at 10 min following CSF1 stimulation (Figure 4A). Rac1 activation is sufficient to drive membrane ruffling and macropinocytic cup formation, but subsequent Rac1 inactivation is required for efficient cup closure and progression toward a mature macropinosome (Yoshida *et al*., 2009; Fujii *et al*., 2013) suggesting that a Rac1 effector needs to be switched off. The persistence of actin-rich ruffles following *Pak2* disruption therefore raises the possibility that PAK2 somehow limits the coordinated plasma membrane remodeling events that accompany cup closure and the transition from an actin-rich plasma-membrane domain to an internalized macropinosome.

**Figure 4.**
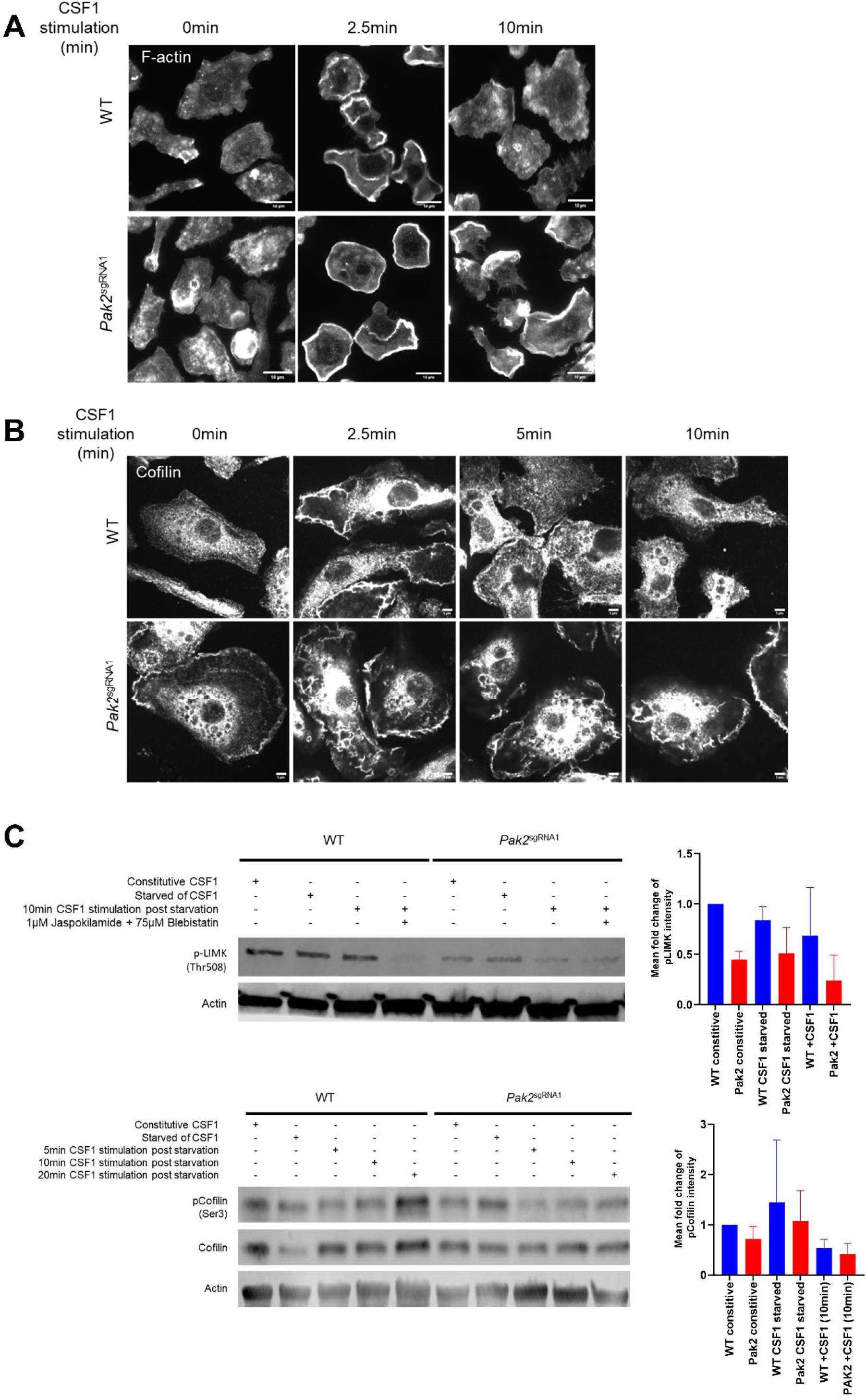
PAK2 regulates actin turnover and LIMK/cofilin signaling during CSF1-induced membrane ruffling. A. *Pak2*^sgRNA^ BMDM has sustained filamentous actin in lamellipodia following growth factor stimulation, suggesting that PAK2 is required for efficient actin turnover during macropinocytosis. BMDM starved of CSF1 overnight were stimulated with 200ng/mL CSF1 and were 4% PFA fixed at indicated times. F-actin stained with Alexa-488 Phalloidin and imaged on 40X air on Olympus IX83. B. Sustained cofilin presence in membrane ruffles and macropinosomes can be seen in *Pak2*^sgRNA^ BMDM relative to WT. Imaged on 60X oil spinning disk confocal microscope. C. PAK2 depletion reduces LIMK1 phosphorylation and alters cofilin regulation. In PAK2^sgRNA^ BMDM, a lower level of phosphorylated LIMK1 (Thr508) was observed, and a lower level of phosphorylated cofilin (Ser3) was also observed. The bar graph represents mean fold change across independent experiments (n=3).

Cofilin regulates actin dynamics by binding and severing F-actin, and its actin-binding activity is inhibited by phosphorylation at Ser3 by LIM kinases (Moriyama *et al*., 1996; Edwards *et al*., 1999). Cofilin was transiently associated with membrane ruffles and macropinosome-associated structures in WT BMDMs, whereas prominent cofilin localization persisted at these structures in *Pak2*^sgRNA^ BMDMs following CSF1 stimulation (Figure 4B). PAK2 depletion was also associated with reduced phosphorylation of LIMK1 at Thr508 and altered cofilin abundance and phosphorylation, including lower phospho-cofilin Ser3 under several conditions (Figure 4C). Together, these results indicate that PAK2 loss disrupts LIMK/cofilin regulation and the normal kinetics of actin remodeling during CSF1-induced macropinocytosis.

*PAK2 depletion alters phosphorylation of proteins associated with cytoskeletal and endosomal regulation* Because PAK2 is a serine/threonine kinase, we used quantitative phosphoproteomic analysis to identify candidate signaling proteins whose phosphorylation was altered following *Pak2* disruption. A subset of phosphoproteins showed decreased abundance in *Pak2*sgRNA BMDMs, including proteins associated with phosphoinositide signaling, cytoskeletal organization, and endosomal trafficking (Figure 5A, Supplemental Table 3). Notable candidates included INPP5D/SHIP1, the Rab5 effector ANKFY1/Rabankyrin-5, EEA1, TBC1D2, CLIP1, and LSP1, suggesting that PAK2-dependent signaling may extend beyond regulation of the actin cytoskeleton to pathways coordinating membrane and endosomal dynamics. These changes could reflect direct PAK2-dependent phosphorylation or indirect effects on downstream kinases, phosphatases, or signaling complexes.

**Figure 5.**
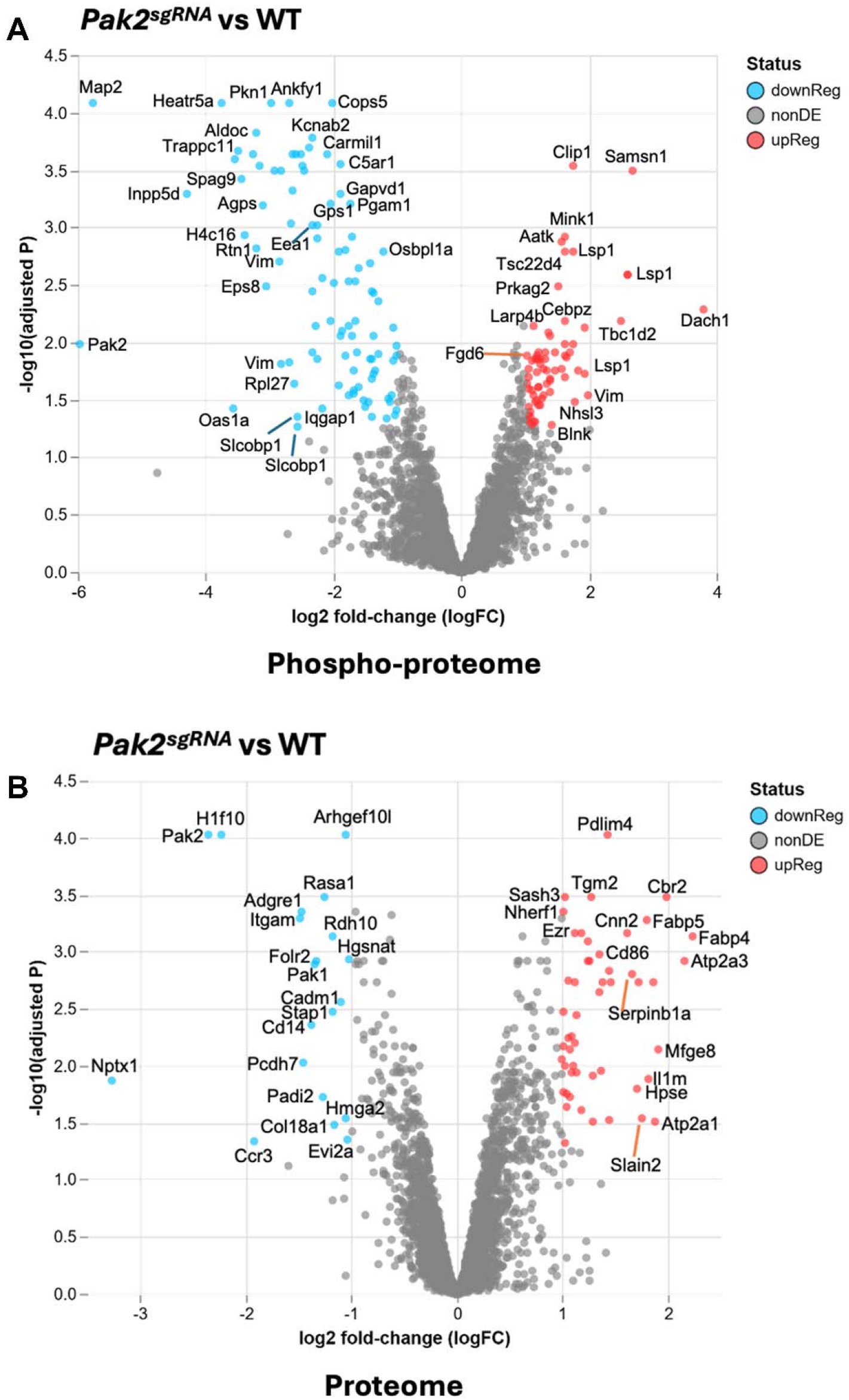
Proteomic and phosphoproteomic analyses identify candidate pathways downstream of PAK2 in BMDMs. A) Volcano plot showing phosphorylated targets in phospho-proteome screen. B) Volcano plot showing targets in proteome screen

Parallel analysis of the total proteome (Supplemental Table 4) showed that, with three exceptions, the proteins highlighted by the phosphoproteomic analysis were not correspondingly decreased in abundance. Thus, the reduced phosphopeptide abundance observed following *Pak2* disruption predominantly reflects changes in phosphorylation rather than simply reduced expression of the corresponding proteins (Figure 5B, Supplemental Table 4). The phosphoproteomic dataset therefore identifies candidate PAK2-dependent signaling pathways that may contribute to the altered cytoskeletal and membrane-trafficking phenotypes of PAK2-depleted macrophages.

## Discussion

Here, genome-wide CRISPR/Cas9 screening in primary BMDMs identified both positive and negative regulators of macrophage fitness and unexpectedly revealed PAK2 as a suppressor of macrophage proliferation. Because sgRNA delivery occurred after several days of CSF1-dependent culture and cell adherence, enrichment of *Pak2*-targeted cells is most consistent with altered fitness of nascent macrophages rather than an effect restricted to granulocyte-macrophage progenitors. Targeted *Pak2* disruption confirmed increased BMDM accumulation and maintained cyclin-D1 expression, while also producing enhanced membrane ruffling, larger macropinosomes, increased fluid-phase uptake, persistent actin-rich structures, altered LIMK/cofilin signaling, and broad changes in the phosphoproteome. Together, these findings identify PAK2 as a context-dependent regulator that restrains macrophage proliferation and membrane-remodeling activity.

The suppressive effect of PAK2 on macrophage proliferation contrasts with its predominantly pro-fitness functions described in several transformed cell types. PAK2 promotes proliferation and transformation in epidermal and melanoma cells through c-Jun-dependent signaling, with PAK2 depletion reducing proliferative and transforming activity (Li *et al*., 2011). PAK2 also supports proliferation in NF2-deficient schwannoma cells, where its depletion reduces Wnt/β-catenin signaling, c-Myc, and cyclin D1, and pharmacologic inhibition of group I PAKs suppresses schwannoma growth and tumorigenesis (Zhou *et al*., 2011; Licciulli *et al*., 2013). PAK2 additionally contributes to tumor-cell invasion, with depletion impairing invasion of breast carcinoma and medulloblastoma cells through mechanisms distinct from PAK1 (Coniglio *et al*., 2008). More recently, PAK2 was identified as a selective dependency in mesenchymal colorectal cancer, where its loss impaired growth and metastatic capacity (Buikhuisen *et al*., 2023). Together, these studies establish a predominantly growth- and tumor-promoting role for PAK2 in several transformed-cell contexts, making the increased proliferation observed after *Pak2* disruption in BMDMs particularly striking.

PAK2 functions as a brake on macrophage membrane remodeling rather than as a simple downstream effector required for Rac-driven ruffling. Macrophages place particularly strong demands on Rac-dependent cytoskeletal signaling because membrane ruffling, phagocytosis, and constitutive or growth factor-stimulated macropinocytosis require repeated spatial and temporal activation of Rho-family GTPases (Wells *et al*., 2004; Redka *et al*., 2018; Doodnauth *et al*., 2019). Consistent with our findings, loss or inhibition of PAK activity in macrophages produces extensive membrane expansion and ruffling together with increased macropinocytic and phagocytic uptake (Rochelle *et al*., 2025). Their results complement our observation that *Pak2* disruption increases macropinosome size and Lucifer yellow uptake, providing independent evidence that PAK2 restrains macrophage engulfment. This phenotype is particularly notable because PAK2 is a canonical effector of Rac1 and Cdc42 (Manser *et al*., 1994). Rac1 signaling during macrophage macropinocytosis, however, is highly dynamic: Rac1 activation accompanies ruffle and cup formation, whereas Rac1 must subsequently be inactivated for efficient cup closure and progression to an internalized macropinosome (Yoshida *et al*., 2009; Fujii *et al*., 2013). Thus, the persistence of robust ruffling and increased macropinocytosis after *Pak2* disruption argues against a simple Rac1-PAK2 pathway in which PAK2 is required to drive actin polymerization. Instead, PAK2 may represent a Rac-responsive pathway that that must be turned off to enable plasma membrane fusogenicity and macropinocytic cup sealing.

The altered phosphoproteome of PAK2-depleted macrophages may reflect both loss of PAK2-dependent signaling and secondary changes produced by remodeling of the macropinocytic and endosomal system. Protein phosphorylation reflects the balance of kinase and phosphatase activities, and the increased number, size, or persistence of macropinosomes following *Pak2* disruption could itself change this balance by redistributing signaling proteins or altering their residence time on endosomal membranes. Among the proteins with reduced phosphorylation, INPP5D/SHIP1 is particularly intriguing because SHIP1 is expressed predominantly in hematopoietic cells and acts as an important negative regulator of PI3K-dependent signaling, proliferation, survival, and cytoskeletal responses in macrophages and other immune cells (Rauh *et al*., 2004; Pauls and Marshall, 2017). Reduced SHIP1 phosphorylation therefore raises the possibility that PAK2 intersects with a hematopoietic-enriched signaling circuit that could contribute to its distinct effects in macrophages. GAPVD1 provides a second mechanistic clue because its VPS9 domain regulates Rab5-dependent early endocytic trafficking, providing a potential link between PAK2 signaling and the transition from plasma-membrane remodeling to endosomal organization (Bean *et al*., 2018).

The opposing effects of PAK2 on tumor-cell and macrophage fitness raise an intriguing possibility that targeting PAK2 could have complementary or synergistic effects on malignant cells and macrophages within the tumor microenvironment. PAK2 has been proposed as a therapeutic vulnerability in cancers in which it supports cytoskeletal remodeling, growth, invasion, and metastatic progression; for example, PAK2 deletion or inhibition markedly impairs growth and metastasis of mesenchymal colorectal cancer models (Buikhuisen *et al*., 2023). PAK2 has been proposed as a therapeutic vulnerability in cancers in which it supports cytoskeletal remodeling, growth, invasion, and metastatic progression; for example, PAK2 deletion or inhibition markedly impairs growth and metastasis of mesenchymal colorectal cancer models (Buikhuisen *et al*., 2023). Conversely, our findings indicate that PAK2 inhibition would not be expected to suppress macrophage fitness in the same manner. Recent work further demonstrates that loss of PAK2 activity in macrophages enhances macropinocytosis and phagocytosis and can increase uptake of cancer cells, in part through altered endosomal trafficking of inhibitory receptors such as SIRPA (Rochelle *et al*., 2025). These observations raise the possibility that PAK2-directed therapies could simultaneously impair PAK2-dependent tumor cells while enhancing selected macrophage uptake functions. Whether such effects are beneficial will depend on tumor context, macrophage state, inhibitor specificity, and the consequences of sustained PAK inhibition *in vivo*. Nevertheless, the strikingly different consequences of PAK2 loss in macrophages and transformed cells underscore the importance of defining cell-type-specific kinase functions when developing targeted therapies.

## Supplemental Movies Legends

**Supplemental Movie 1:** Brightfield microscopic movie of WT BMDMs related to Figure 3C. Cells were starved overnight in DMEM containing 10% FBS and stimulated with CSF1. Cells were imaged in HBSS at 37°C using an Olympus IX83 microscope platform (Olympus, Shinjuku City, Tokyo, Japan) with a 60× oil-immersion objective (NA 1.42). Brightfield images were acquired every 15 s using a cooled CCD camera for a total duration of 5 min. Scale bar = 5 μm.

**Supplemental Movie 2:** Brightfield microscopic movie of *Pak2*^*sgRNA*^ BMDMs related to Figure 3C. Cells were starved overnight in DMEM containing 10% FBS and stimulated with CSF1. Cells were imaged in HBSS at 37°C using an Olympus IX83 microscope platform (Olympus, Shinjuku City, Tokyo, Japan) with a 60× oil-immersion objective (NA 1.42). Brightfield images were acquired every 15 s using a cooled CCD camera for a total duration of 5 min. Scale bar = 5 μm.

**Supplemental Movie 3:** Brightfield microscopic movie of Non-target BMDMs related to Figure 3C. Cells were starved overnight in DMEM containing 10% FBS and stimulated with CSF1. Cells were imaged in HBSS at 37°C using an Olympus IX83 microscope platform (Olympus, Shinjuku City, Tokyo, Japan) with a 60× oil-immersion objective (NA 1.42). Brightfield images were acquired every 15 s using a cooled CCD camera for a total duration of 5 min. Scale bar = 5 μm.

## Acknowledgements and Funding

Research reported in this publication is supported by NIH grants R15GM139162, R35GM131720, R01GM157516, P20GM135008. Content is solely the responsibility of authors. This material is based on work conducted using SDSU Genomics Sequencing Facility (RRID:SCR_023959) & Functional Genomics (RRID:SCR_023786) facilities, supported in part by NSF/EPSCoR Grants 0091948 & IIA-1355423, SD AES, and by the State of South Dakota. Any opinions, findings, and conclusions or recommendations expressed in this material are those of the authors and do not necessarily reflect the views of the National Science Foundation (USA).

