## Supplemental Figure 1-2 for "High-resolution CRISPR/Cas9 screens identify PAK2 as a suppressor of macrophage proliferation"

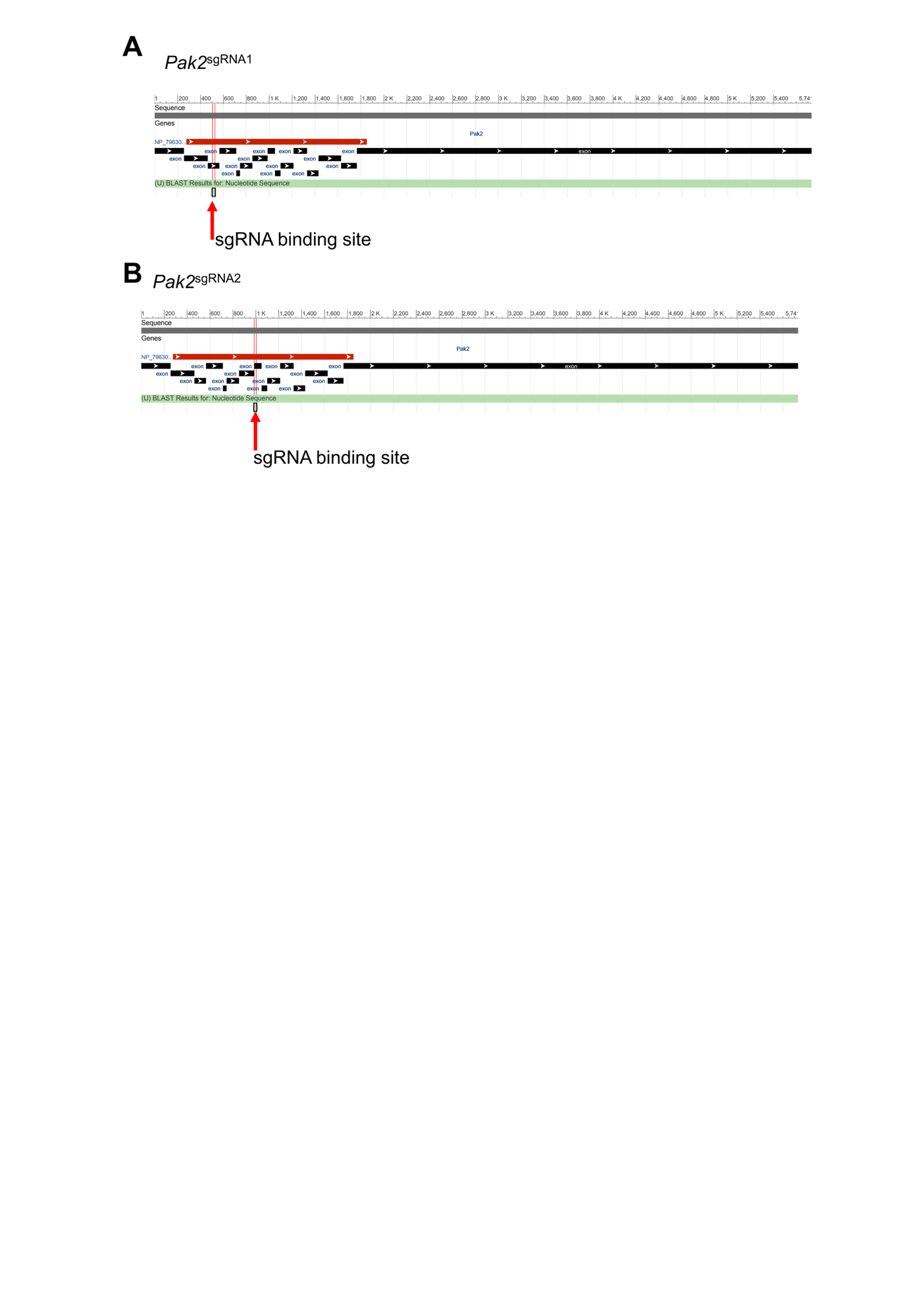

**Supplemental Figure 1. sgRNA binding sites on mouse genome.** sgRNA designed to target *Pak2* were aligned to the mouse genome using NCBI Blast tool. sgRNA1 is complementary to a region on exon 3 and sgRNA2 is complementary to a region in exon 8.

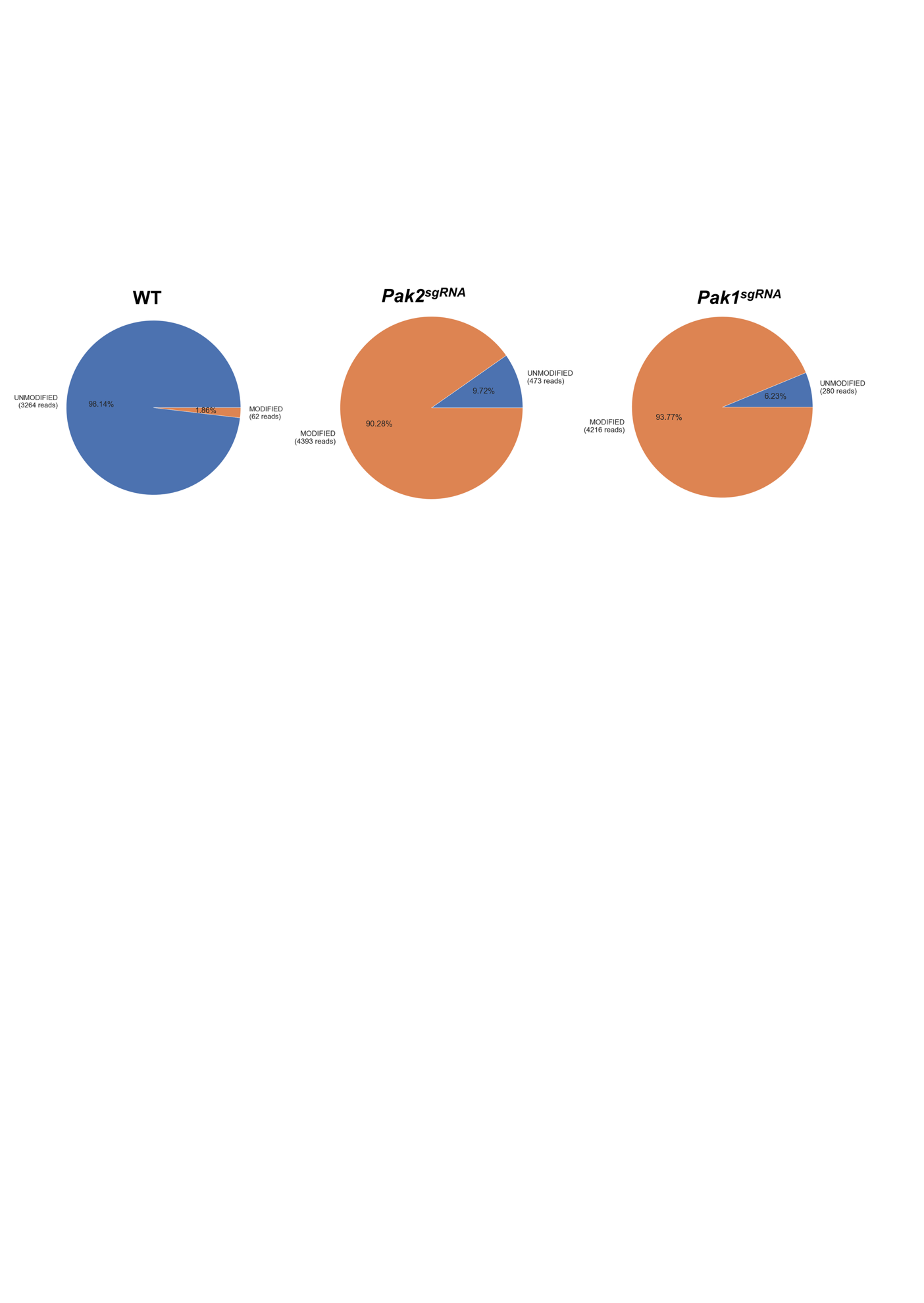

**Supplemental Figure 2. Cut-site sequencing for Pak1 and Pak2 disruption.** Nanopore sequencing was used to confirm the disruption of target genes at the location targeted by sgRNA. Pie charts show the distribution of modified and unmodified alleles from lysates of BMDM transduced with the indicated sgRNA.
